# SMA Type II Skeletal Muscle Treated with Nusinersen shows SMN Restoration but Mitochondrial Deficiency

**DOI:** 10.1101/2024.02.29.582680

**Authors:** Fiorella Carla Grandi, Stephanie Astord, Sonia Pezet, Elèna Gidaja, Sabrina Mazzucchi, Maud Chapart, Stéphane Vasseur, Kamel Mamchaoui, Piera Smeriglio

**Affiliations:** Sorbonne Université, INSERM, Institut de Myologie, Centre de recherche en Myologie F-75013 Paris, France; Centre de Ressources Biologiques - Myobank-AFM de l’Institut de Myologie. Hôpital de la Pitié-Salpêtrière F-75013 Paris, France

**Keywords:** Spinal Muscular Atrophy, transcriptomics, muscle, SMN1, mitochondria

## Abstract

Spinal muscular atrophy (SMA) is a rare autosomal recessive developmental disorder caused by the genetic loss or mutation of the gene *SMN1 (Survival of Spinal Motor Neuron 1)*. SMA is classically characterized by neuromuscular symptoms, including muscular atrophy, weakness of the proximal muscles, especially those of the lower extremities, and hypotonia. Although originally thought of as a purely motor neuron disease, current research has shown that most, if not all, tissues are affected, including the muscle. Until recently, muscle problems in SMA were predominantly considered a consequence of denervation due to the motor neuron death. However, recent work using muscle-specific mouse models of SMN loss, as well as skeletal stem cell specific models have shown that there are tissue specific problems in muscle due to SMN deficiency. Several years ago, SMA treatment underwent a radical transformation, with the approval of three different SMN*-*dependent disease modifying therapies. This includes two *SMN2* splicing therapies - Risdiplam and Nusinersen, which can be administered by Type II patients that have symptom onset later in age. One main challenge for Type II SMA patients treated with Risdiplam and Nusinersen is ongoing muscle fatigue, limited mobility, and other skeletal problems, including hip dysplasia and scoliosis. To date, few molecular studies have been conducted on SMA-patient derived tissues after treatment, limiting our understanding how different organ systems react to the therapies, and what additional combination therapies may be beneficial. With this goal in mind, we collected paravertebral muscle from the surgical discard in a cohort of 8 SMA Type II patients undergoing spinal surgery for scoliosis, as well as 7 non-SMA controls with scoliosis and used RNA-sequencing to characterize their molecular profiles. We observed that despite a restoration of the SMN mRNA and protein levels in these patients – at levels at or above the controls – a subset of patients continued to have alterations in mitochondrial metabolism and other markers of cellular stress.

## INTRODUCTION

Spinal muscular atrophy (SMA) is a rare autosomal recessive developmental disorder caused by the genetic loss or mutation of the gene *SMN1 (Survival of Motor Neuron 1)* ^1^. The disorder is classically characterized by neuromuscular symptoms, including muscular atrophy, weakness of the proximal muscles, and hypotonia^1,2^. At the cellular level, *SMN1* loss results in the death of motor neurons (MN) which innervate the muscle. At the molecular level, SMN has a variety of associated molecular functions, including RNA splicing, R-loop resolution, and cytoskeletal dynamics^3^.

SMA symptoms exists on a spectrum and patients are classified in five stratifications (SMA Type 0 – 4) which guide clinical care^2,4^. These groups are determined based on the degree of motor symptom involvement and the age of onset. For example, Type 0 SMA is prenatally lethal while Type 4 SMA is diagnosed in adulthood and can exhibit muscle weakness and limited mobility. Additionally, patients are stratified by the number of copies of the gene homologue *SMN2*, which is the major genetic modifier of the disease. *SMN2* is a nearly homologous copy of *SMN*, situated in the same 5q13 locus which has undergone events of duplication and deletion. Therefore, *SMN2* can be found in varying copy numbers in the human genome^5,6^. However, *SMN2* differs from *SMN1* by a few bases that modify a splice junction, inducing a very low production of the full-length SMN (10%) and 90% transcription of a truncated SMN mRNA without exon 7, classically referred to as *SMNdelta7*. This transcript is unstable, has a shorter half-life, and cannot fully compensate for the function of *SMN1*. The critical importance of the dose of the SMN protein is underscored by the relative amelioration of symptoms in type II-IV SMA patients, which have increasing numbers of SMN2 – Type I patients usually have 2 copies of *SMN2* while Type III patients have 3-4 copies^2^.

Classically, SMA has been considered a disease of motor neurons. However, as *SMN* is ubiquitously expressed through the body^7,8^ the tissue-specificity of this effect has been hard to understand. Emerging work has demonstrated that SMA is a whole-body developmental disorder, with SMN exerting specific roles in several tissues, including the pancreas, mesenchymal tissues, liver, brain and muscle^9–11^. Thus, understanding the role of SMN in non-MN tissues is necessary for clinical management. In particular, work from several groups has highlighted cell-intrinsic defects in muscle tissue in mouse models where *Smn1* is lost only in one or several muscle cell types but not in motor neurons^11,12^. This work strongly suggests that restoring SMN to the muscle is critical for optimal care.

Several years ago, SMA treatment underwent a radical transformation, with the approval of three different *SMN-*dependent disease modifying therapies^1^ which revolutionized the life expectancy of patients. However, these treatments remain non-curative. Of these, the most efficacious is the gene therapy Onasemnogene abeparvovec (ZOLGENSMA)^13^, which is available as a treatment for infants with Type I SMA, rendering this previous fatal disease survivable. Long-term follow-up of treated patients has demonstrated significant gain in motor function and motor milestones, including sitting, crawling and head control^13^. For those patients who are not eligible for ZOLGENSMA treatment – either because of age-of-symptom onset, current age, or AAV immunoreactivity - two other treatments are available which act on *SMN2*^1^ splicing mechanism, Nusinersen (Spinraza) and Risdiplam (Evrysdi). Both Nusinersen and Risdiplam are SMN-dependent therapies whose mechanism of action is based on inducing the inclusion of exon 7 in the remaining *SMN2* copies^1,14,15^. Nusinersen is an antisense oligonucleotide (ASO), which modulates the splicing site of *SMN2* and enhances the translation of the fully functional SMN protein^16^. It is injected intrathecally, and treatment must be administered repeatedly. The first loading doses are typically administered at 14-day intervals, with a 4^th^ loading dose administered 30 days after the 3^rd^ dose. Finally, maintenance doses are administered once every 4 months afterwards. Risdiplam is the only oral SMN-targeted therapy, consisting of a small-molecule pre-mRNA splicing modifier that also targets exon 7 splicing^17,18^. The drug can be administered daily. The effectiveness of these therapies, which were originally designed to target the central nervous system, in peripheral tissues remains poorly understood.

While these treatments have been life changing for SMA patients, the road ahead contains challenges for patients and clinicians since the three treatments do not always lead to a full restoration or alteration of clinical symptoms. The clinical trial data from Nusinersen showed a wide range of clinical efficacy, with some patients attaining motor function improvement, while others had little changes, especially among adult patients^14,19–21^. In part, this is due to the need for early treatment, putatively before symptom onset, to fully restore all SMN-related deficiency problems^22,23^. This may be partially because the levels of SMN are highest prenatally^24^. However, in older children and adult with SMA, which collectively represent two-thirds of the SMA population, early gene therapy treatment is not an option.

Approximately 10,000 SMA patients have been treated with Nusinersen, however, this suggests that new phenotypes, previously not seen due to the fatality of the disease, may emerge and may need additional SMN-independent therapies.

Type 2 SMA patients represent about 20% of all cases and have an onset of symptoms between 6 and 18 months, with an unassisted life expectancy of 25 years, which has been vastly improved by the new therapies and supportive care. Most patients achieve sitting milestones, although sometimes with delay, and by clinical definition, these patients generally do not sit or walk interpedently. They display proximal predominant weakness, especially in the lower limbs, and most, if not all, have scoliosis^25^. Due to the age of onset, clinical management includes treatment with Risdiplam and Nusinersen injections. However, post-treatment challenges remain, including muscle fatigue, limited mobility, and other skeletal problems, including such as hip dysplasia and scoliosis^26^.

To date, few molecular studies have been conducted on SMA patient tissues after treatment^24^, limiting our understanding on how different organ systems react to the therapies, and what additional combination therapies may be beneficial. State-of-the-art SMA therapies directly seek to increase the amount of SMN protein. Molecular studies on the muscle of the treated patients are lacking. A previous work on *SMN* mRNA splicing in several tissues following ASO injection had found induction of proper splicing only in the spinal cord but not in the brain or other peripheral tissues of treated SMA patients, including the iliopsoas and diaphragm muscles ^24^. However, that study was based on post-mortem samples from young children (<72 months), with two copies of *SMN2*, consistent with an SMA Type I profile ^2,24^. Many of these samples were procured from severe cases not long after the first injection. As treatment with Nusinersen is currently mostly used in Type II SMA patients or in adults, we wanted to characterize the result of treatment in this patient group. Therefore, we performed a histopathological and molecular characterization of paravertebral muscle from the surgical discard in a cohort of 8 treated SMA Type II patients undergoing spinal surgery for scoliosis as well as 7 non-SMA controls. We observed that despite a restoration of the SMN mRNA and protein levels in these patients – at levels at or above the controls – a subset of patients continued to have alterations in mitochondrial metabolism and other markers of cellular stress. We observed a marked heterogeneity in the muscle phenotype and expression landscape and this likely corresponds to a variable patient’s response to treatment. This analysis brings new understandings on the peripheral effect of SMN splicing therapies and confirms the importance of combined SMN dependent and independent treatments to maximize the skeletal muscle restoration in type II patients.

## MATERIALS AND METHODS

### Ethical consent for study of human samples

Samples were obtained with patient or parental consent, collected from the surgical residue from patients undergoing surgery for spinal sclerosis. Samples were collected under the authorization of the minister of research approval number AC-2019-3502, in accordance with French and European laws.

### Sex as a biological variable

In this study, biological sex was taken into consideration. Our study sought to examine a sex balanced cohort. In the SMA cohort, we had a sex balanced group, with 4 female and 4 male samples. However, due limited sample availability with the controls, we have a 5 female and 2 male control samples. We explicitly tested for differences in the SMA presentation based on sex, but no correlation was found between SMA subgroups at sex, and no sex-effect was found among the differentially expressed genes.

### Mouse husbandry

Animals were bred and housed as previously described^27^ under controlled conditions following the French and European guidelines for the use of animal models (2010/63/EU). Briefly, breeding pairs of triple transgenic (*SMN2^+/+^, SMNΔ7*^+/+^, *Smn*^+/−^; no. SN 5025) were purchased from The Jackson Laboratory. The genotype of the offspring can be WT (*SMN2*^+/+^, *SMNΔ7*^+/+^, *Smn*^+/+^), heterozygous (*SMN2*^+/+^, *SMNΔ7*^+/+^, *Smn*^+/−^), or knockout, named SMNΔ7 (*SMN2*^+/+^, *SMNΔ7*^+/+^, *Smn*^−/−^). Mice were genotyped using the previously established primers. Mice were euthanized 14 days post-injection for SMN expression analyses by intraperitoneal anesthetic injection (10 mg/kg xylazine, 100 mg/kg ketamine), followed by intracardiac perfusion, with PBS. Euthanasia was performed according to regulations and muscle samples were recovered and immediately frozen in liquid nitrogen for further downstream processing.

### DNA extraction from human muscle samples

DNA was extracted from ∼2-3mg of frozen muscle tissue during the Qiagen DNeasy Blood and Tissue Kit. Samples were first ground into a powered using a tissue grinder and then digested for 3-6 hours with proteinase K as per the manufacture’s recommendations.

### Mitochondrial DNA (mtDNA) copy number calculation

Mitochondrial copy number was calculated from the extracted DNA using a previously validated protocol^28^. Briefly, DNA was diluted to 10ng<u>/µl</u> in water, and 2 µl were loaded into a 25µl SYBR Green reaction, as per the protocol. Samples were normalized using a genomic DNA housekeeping gene.

### RNA extraction from human muscle samples

RNA was extracted using the Qiagen RNAeasy Mini Kit, with some modifications. Frozen samples (∼3-4 mg) were ground in a tissue grinder, making sure to keep the tissue frozen. The samples were then incubated with 800ul of RLT buffer and passed through a 20G needles several times until the homogenized lysate was no longer viscous. Samples were then processed according to the manufacture’s specifications. RNA quality was assessed using an Agilent TapeStation. All samples had a RIN > 6. For cells, 500,000 cells were collected by spinning and resuspended in RLT buffer according to the manufacturer’s specifications.

### Real-time quantitative PCR for SMN1/2 expression

From the RNA extracted from different cell lines, 1µg was converted into cDNA using the High-Capacity cDNA Reverse Transcription Kit (Applied Biosciences) according to the manufacture’s recommendation. 10µll PCR reactions were set up in duplicate using the TaqMan Mix (Applied Biosciences) with SMN1/2 probe Hs00165806_m1 and HPRT as a housekeeping control. Relative expression was calculated using the delta delta CT method.

### cDNA library generation and sequencing

Libraries for sequencing were prepared using the SMART-Seq® v4 PLUS Kit (Takara/CloneTech) according to the manufacture’s specifications, using 10 nanograms of input RNA. Samples were sequenced after barcoding using the Illumina NovaX platform with 150 bp paired-end reads.

### Library alignment and differential gene expression analysis

Fastq files were analyzed using the nf-core RNA seq pipeline with the standard parameters^29^. Reads were aligned to hg38. Differential expression analysis was performed using DESeq2^30^, and R was used for visualization of the data. Pathway analysis was performed using EnrichR^31^ and Metascape^32^.

### Fiber Type Deconvolution from RNA-sequencing data

To determine the predicted percentage of each fiber type in our RNA-sequencing data, we utilized the previously published profiles from Oskolkov et al^33^.

### Hemotoxin and eosin staining

12 µm sections were obtained from each sample using a cryostat (Leica) and fixed in 4% PFA for 10 min. Slides were immersed in PBS for 2 min. Tissues sections were incubated in Mayer hematoxylin (Biognost, #HEMH-OT-500) staining solution for 3 minutes and then in running tap water for 5 minutes followed by 30 seconds of incubation in Eosin Y (ScyTek, #EYQ500), Slides were rinsed in running tap water and then dehydrated in alcohol and xylene. Slides were mounted using VectaMount™ Mounting Medium (Vector Laboratories). Images were obtained using a digital microscope (Keyence, VHX-7000)

### Sirius Red

Slides were washed 5 minutes with tap water and then put 5 minutes in 100% ethanol before letting them dry for 20 minutes. Slides were incubated in Sirius Red (Sigma, #365548 – 0.3g in 100ml of picric acid) for 1 hour and then in running tap water (5 minutes) and acetic acid (0,5%, 5 minutes) followed by dehydration and mounting with VectaMount™ Mounting Medium (Vector Laboratories). Images were obtained using a digital microscope (Keyence, VHX-7000) and quantification realised using ImageJ software. All histological analyses were performed in a blinded manner.

### Fiber size measurement

Image J^34^ was used to quantify the area in arbitrary units. Images with intact myofibers were selected, all taken at the same Z100: X200 objective zoom on the Keyence VHX-7000. Individual whole fibers were traced by hand and quantified using the ‘measure’ area function. All histological analyses were performed in a blinded manner.

### Total cholesterol and cholesterol-ester measurements from muscle samples

Total cholesterol and cholesterol esters were measured using the Cholesterol Ester-Glo Assay (Promega cat # J3190) using the manufacturer’s instructions. Tissue lysate (1:25) was obatined using around 15mg of tissue disrupted in 500µl of lysis solution with a homogenizer.

### Protein extraction and western blotting

Protein extracts were prepared from liquid nitrogen frozen human or muscle tissues. Tissues were lyzed in (RIPA buffer (Sigma) supplemented with a protease inhibitor cocktail (Complete Mini, Roche Diagnostics). Proteins were run on 4-10% Bis-Tris gels (BioRaD) and transferred membranes using the TurboTransfer System (BioRad). The membranes were then incubated with an anti-SMN antibody (1:1,000; BD Biosciences), anti-vinculin (1:1000; Sigma Cat V9191), anti-phospho (Ser15)-p53 (1:1000, Cell Signaling Cat#92845), or anti-GADD45A (1:1000, Santa Cruz Cat # /sc-6850) diluted in TBST blocking buffer (Tris-buffered saline containing 0.2% Tween 20, BioRad) supplemented with 5% non-fat dry milk. Primary antibody incubation was performed overnight. After 3 washes in TBST buffer, the membranes were incubated with a horseradish peroxidase-conjugated against the appropriate species antibody (1:10,000; GE Healthcare) diluted in the blocking buffer. The membranes were further revealed using the chemiluminescence substrate SuperSignal Ultra reagent (Pierce) and image on a BioRad ChemiDoc MP Imaging System.

### miRNA quantification assay by real-time PCR

miRNA quantification was performed from total RNA extractions using the TaqMan MicroRNA Reverse Transcription Kit (Applied Biosciences) according to the manufacturer’s instructions. Briefly, samples were retrotranscribed using the specific RT probe for each miRNA (Applied Biosciences: miR-1 2222 mir-206 000510 miR-24 402 U6 1973). Afterwards each sample was quantified using a specific primer-probe set. Quantification was performed according to the delta-delta Ct method, using U6 as the housekeeping gene.

### Myoblast derivation, culture, and differentiation

Immortalized myoblasts from control (AB1190; male 16-years old and KM1421, female 13 years old) and SMA (KM432 SMA I bis11PV and KM1150SMAII7PV) paravertebral muscle were obtained from the MyoLine Platform. Cells were cultured at 37C at 5% CO_2_ under humidity in myoblast proliferation medium [1Vol medium 199 (ThermoFisher, 41150020) + 4 vol DMEM (ThermoFisher, 61965-026) + 20% FBS (Gibco) + Gentamycin (50 μg/ml; ThermoFisher, 15750-045) supplemented by Fetuin (25μg/ml; LifeTechnolgoies 10344026), hEGF (5ng/ml, Life Technologies: PHG0311), bFGF(0,5ng/ml; LifeTechnologies PHG0026), Insulin(5μg/ml, Sigma 91077C-1G), Dexamethasone (0,2μg/ml, Sigma: D4902). Cells were passaged with 0.05% Trypsin (ThermoFisher) and routinely tested for mycoplasma (Lonza). For any assays performed in the ‘myoblast’ stage, cells were taken in the proliferation phase (∼70-80% confluent). For differentiation assays (myotubes), 24-well plates were coated with Matrigel (Corning) diluted at 1:20 in media. 150,000 cells were plated and allowed to attach for 2-5 hours, resulting in a confluent plate. The plate was then rinsed with PBS twice, and cells were changed into differentiation media (DMEM + 10 μg/ml of insulin (+Gentamycin 50 μg/ml)). Media was change at 50% volume every 2-days during the seven-day differentiation protocol.

### Statistics

All statistical analysis for RNA, protein and image quantifications were performed using Prism 10 using the test specified in the legend. For RNA-sequencing, statistical testing was done using DeSEQ2 with standard parameters (Fold change 1.4 .5, p.adj <=0.05 and log fold change of the standard error <0.7)). Reported genes are those that passed the adjusted p-value cutoff for multiple hypothesis testing.

## RESULTS

### Treatment with Nusinersen and Risdiplam restores SMN levels in Type II SMA muscle

To study the response of SMA Type II muscle to SMN-splicing therapies, we collected a cohort of paravertebral muscle samples from patients undergoing orthopedic surgery for scoliosis. Our cohort of samples consisted of eight Type II (or Type I bis) SMA patients and 7 non-SMA controls, also undergoing surgery for scoliosis (**Figure 1A**, **Table 1**). Muscle tissue was taken from the surgical discard created during the surgery. To the extent possible, the cohort was sex (9 female, 6 male) and age matched, although on average, the SMA cohort is 4-years younger (mean 12.75 +/- 2 years) than the control cohort (16.14 +/- 1.3 years) (**Figure 1 B**). All SMA patients received intrathecal Nusinersen injections, and a subset had a clinical history of Risdiplam treatment (**Table 1**).

**Figure 1:**
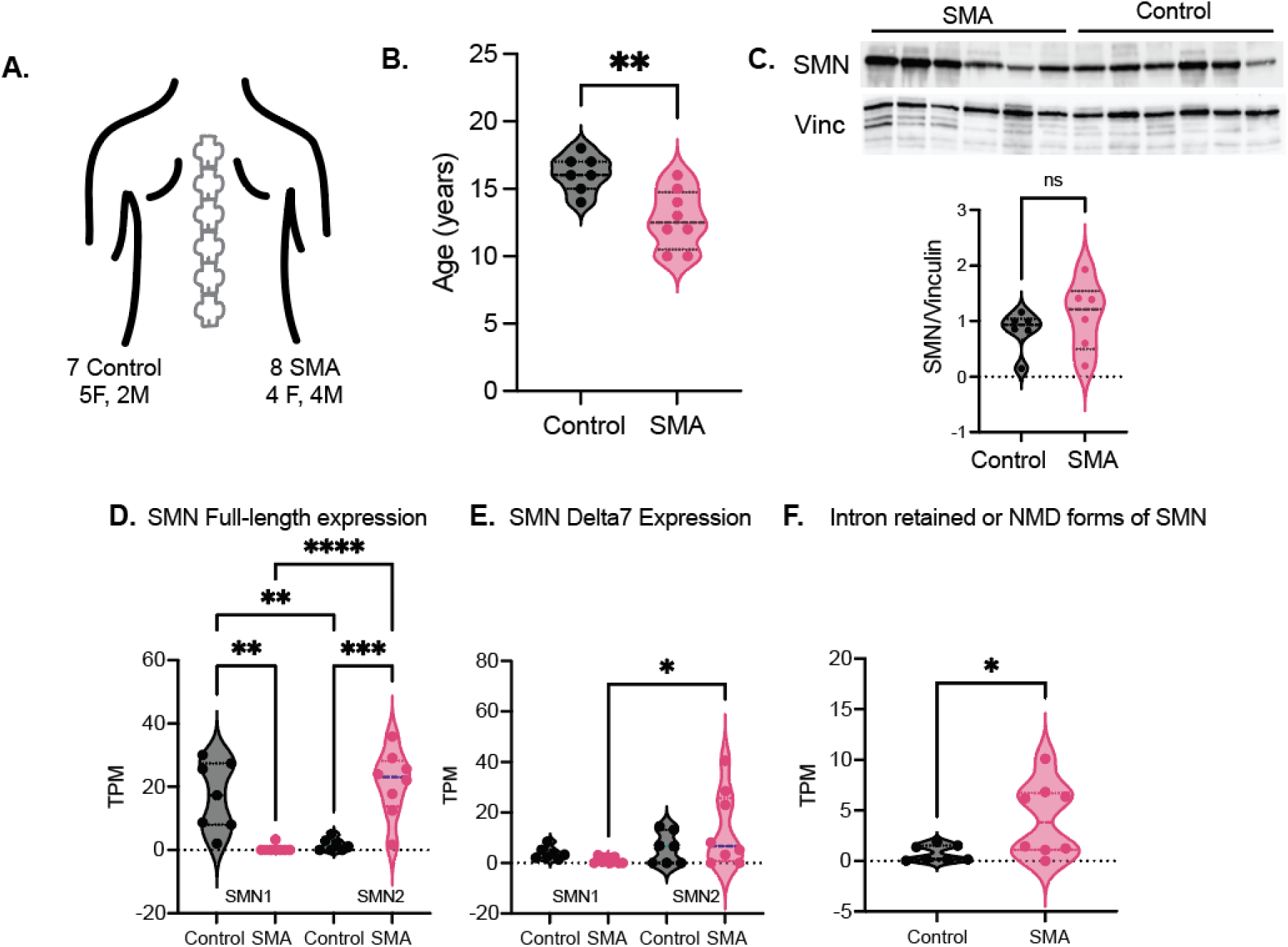
Treatment with Nusinersen and Risdiplam restores SMN protein and RNA levels in Type II SMA paravertebral muscle. **A.** Scheme of the cohort of control and SMA muscle samples, with sex and age balance for each group. **B.** Age, in years, for each group. Groups were compared with a student’s t-test. ** p-value <0.001. **C.** Western blot for SMN protein. Quantification of the blot in the histogram below. Samples were normalized to vinculin. Groups were compared with student’s t-test. **D-F.** Reads, represented as transcripts per million (TPM) mapping to the SMN full length transcript (D), the SMN delta 7 transcript missing exon 7 (E) or any transcript containing retained introns (F). The origin of full-length transcript, either the *SMN1* or *SMN2* locus, is designated below each pair of violin plots. Each point represents a sample. P-values derive from DeSEQ2 differential expression analysis and represent adjusted p-values for multiple hypothesis correction. ** <0.001, ***< 0.0001 and **** <0.00001.

**Table 1:** SMA Paravertebral Muscle Cohort. Table showing the clinical data available for each sample in the cohort. Designations of ‘likely’ indicates we have other samples normally taken at the time of Nusinersen injection but no confirmation from managing clinician.

| Sample ID | Diagnosis | Sex | Age (yrs) | Muscle Type | Treatment notes |
| --- | --- | --- | --- | --- | --- |
| Control 1 |  | F | 15 | Dorsal |  |
| Control 2 |  | F | 18 | Paravertebral |  |
| Control 3 |  | F | 16 | Dorsal |  |
| Control 4 |  | F | 17 | Lumbar |  |
| Control 5 |  | F | 14 | Dorsal |  |
| Control 6 |  | M | 17 | Paravertebral |  |
| Control 7 |  | M | 16 | Dorsal |  |
| SMA1 | SMA Type II | F | 12 | Paravertebral | Risdiplam |
| SMA2 | SMA Type II | F | 10 | Paravertebral | N/A; Nusinersen likely |
| SMA3 | SMA Type II | M | 15 | Paravertebral | Nusinersen |
| SMA4 | SMA Type II | M | 13 | Paravertebral | Nusinersen; Risdiplam |
| SMA5 | SMA Type Ia | F | 16 | Dorsal | Nusinersen |
| SMA6 | SMA Type II | M | 12 | Paravertebral | Risdiplam; likely Nusinersen |
| SMA7 | SMA Type II | F | 10 | Paravertebral | N/A; Nusinersen likely |
| SMA8 | SMA Type II/III | M | 14 | Dorsal | Nusinersen; Risdiplam |

To determine the level of SMN protein – derived either from the *SMN1* or *SMN2* genes - compared between control and SMA patients, we quantified total SMN protein using western blotting. On average, SMA patients had similar or higher levels of SMN protein than controls (**Figure 1C**), suggesting that the splicing based therapies were effective in increasing the amount of SMN in the muscle. To validate these findings at the RNA-level, we performed RNA-sequencing of the same muscle samples. RNA was extracted, cDNA libraries were generated for sequencing, and data was analyzed using the nf-core pipeline^29^ and DeSEQ2^30^.

As total SMN expression, coming from either *SMN1* or *SMN2* copies, is the major disease modifier of severity in SMA^35^, we specifically sought to understand the expression of the various forms of the *SMN* transcript. Although highly similar, *SMN2* contains a change in a splicing acceptor site which makes the propensity of making the *SMNΔ7* transcript more abundant. This transcript generates an unstable protein that cannot fully compensate for the functional full-length SMN protein^6^. Studies have also revealed the presence of an *SMNΔ5* version^36^, although the precise function of this transcript is unknown. Using a variety of SNPs in the introns and UTR regions of the two SMN copies^37,38^, we mapped the reads and assigned them to transcripts annotated from either the *SMN1* or *SMN2* genes. As expected, when comparing the full-length canonical SMN transcripts, we observed a dramatic decrease between control and SMA samples, where few to no reads were mapping to *SMN1*, consistent with the SMA diagnosis (**Figure 1 D**). In control samples, we observed few full-length SMN reads mapping to *SMN2*, with the plurality of *SMN2* mapping reads resulting in the *SMNΔ7* transcript (**Figure 1D & E**). However, in the SMA patients, we observed many more full length than *SMNΔ7* transcripts deriving from the *SMN2* locus (**Figure 1D & E**), in agreement with the mechanism of action of Nusinersen which changes the splicing pattern of *SMN2* to favor the full-length transcript^15^. No correlation was observed between the amount of *SMN2* canonical transcript and the amount of *SMNΔ7* transcripts (**Supplemental Figure 1A**) – rather it 3/8 SMA samples had high *SMNΔ7* levels, irrespective of the amount of *SMN2* full length transcript. We also tested the level of RNA expression with age, as SMN levels have been shown to decrease in aging ^24^, however, within our limited age range, no decrease was observed (**Supplemental Figure 1B**). Consistent with the role of SMN in splicing, we observed a higher degree of SMN transcripts with retained introns in the SMA samples compared to controls (**Figure 1F**). To validate the SMN mapping approach, we used the same pipeline to map reads from a various RNA-sequencing study that profiled the biceps muscle of DMD and SMA Type I patients^39,40^ . We observed few-to-no reads coming from the *SMN1* locus in SMA Type I patients (**Supplemental Figure 1C)** and an increase in *SMNΔ7* transcript from the *SMN2* locus (**Supplemental Figure 1C**).

However, as these patients were not treated with Nusinersen, there was no significant increase in full-length splicing from the SMN2 allele (**Supplemental Figure 1C**). Previous studies have suggested that the amount of SMN full length transcript is also dependent on the levels of two splicing factors, hnRNPA1 and hnRNPA2^41,42^. Our RNA-sequencing data confirm that global expression levels of *HNRNPA1* and *HNRNPA2B1* correlate with increased levels of total SMNΔ7 observed (**Supplemental Figure 1D**). Collectively, this data showed that in Type II SMA patients treated for an extended period, SMN protein and RNA levels are restored in the paravertebral muscle.

### SMA muscles have altered myofiber size with presence of multiple internalized nuclei in a single fiber

We next sought to correlate our molecular phenotypes with a histological analysis of the muscle architecture. We were able to generate high-quality images from 5 out of the 8 SMA samples, potentially due to problems with cryopreservation or storage that compromised the sample architecture in the others. H&E staining demonstrated several classic signs of atrophy and SMA muscle histology, including the presence of hypertrophic fibers (**Figure 2A**) and the clusters of small fibers suggestive of denervation (**Supplemental Figure 2A**). We quantified the area of each myofiber and found that the maximum fiber size, as well as the range of means was increased in the SMA samples (**Figure 2B-C**).

**Figure 2:**
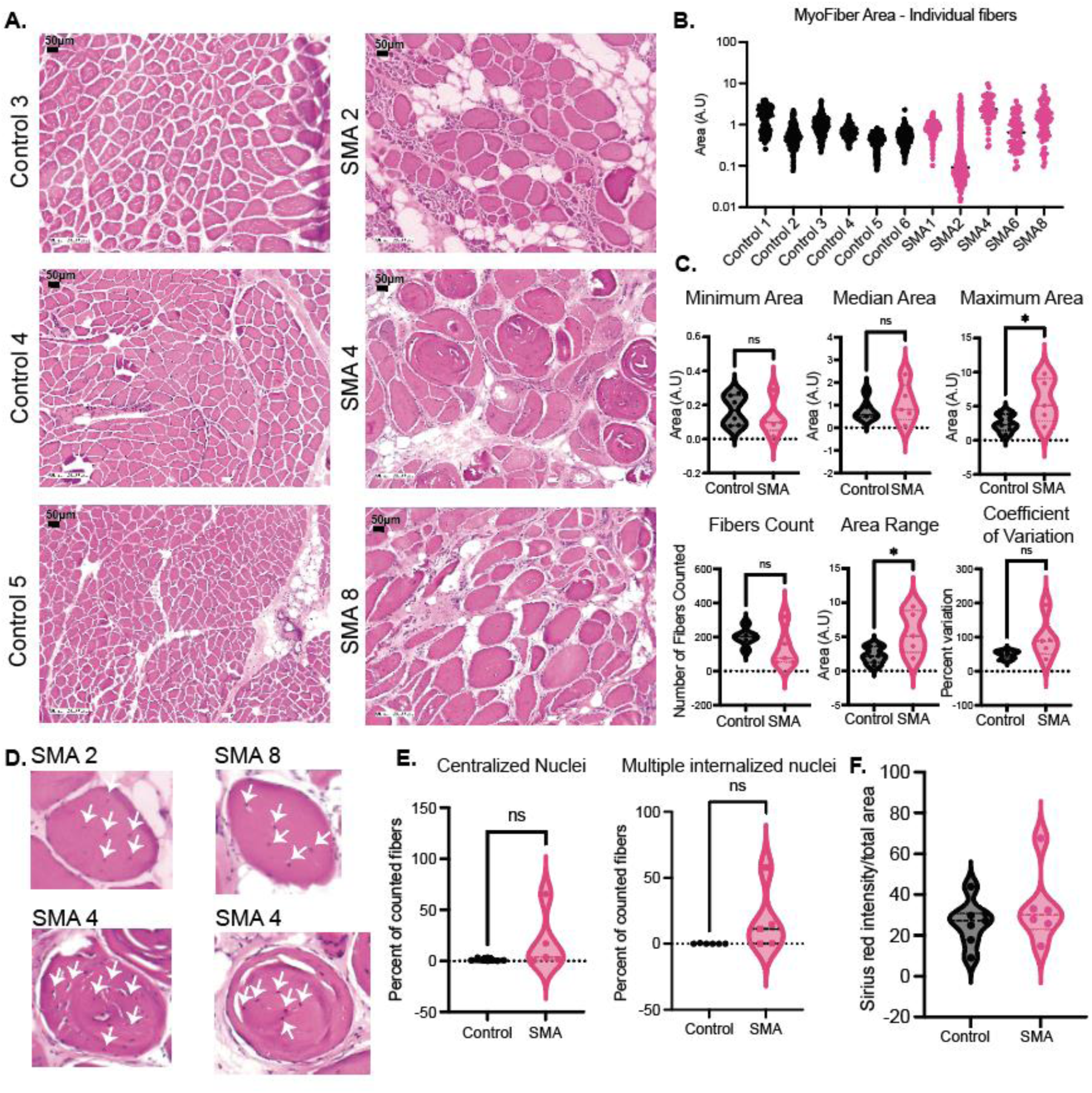
SMA Type II paravertebral muscle after treatment is characterized by abnormal myofiber size distribution. **A.** Hematoxylin and eosin staining of muscle tissues from control and SMA patient samples. Representative images (Z100 x 200) are shown for each group. **B.** Myofiber area for each patient. Each dot represents the area of a single myofiber. **C.** Various summary data of the area measurements from panel B. Each point represents a patient. Samples were compared using the student’s t-test. **D.** Cropped and enlarged myofibers from images showing myofibers with multiple internalized nuclei from the three patients where we observed this phenomenon (P2, P4, P8). Internalized nuclei are designated by white arrows. **E.** Quantification of internalized (both single and multiple) and only multiple centralized nuclei in SMA and controls. Each point represents the percentage of internalized nuclei in the fibers counted in one patient sample. **F.** Quantification of Sirius Red staining for fibrosis detection on control and SMA samples. Groups were compared using the student’s t-test.

Next, we assessed the percentage of the myofibers with internalized nuclei, a common morphological feature of several myopathies^43^. Centralization of nuclei occurs as a response to injury during satellite cells fusion in fiber regeneration. Thus, observing centralized/internalized nuclei in a myopathy can imply that the tissue is attempting to repair itself. We first scored the slides for fibers with internalized nuclei and observed an increase in some but not all SMA samples. In some samples, particular in SMA samples 2, 4 and 8 we observed several myofibers with multiple internalized nuclei (**Figure 2D-E**). However, this was not present in all samples. Within a single tissue section, we could observe both fibers with and without these multiple internalized nuclei in the same cluster, and in general, these multi-centralized nuclei were found in the largest fibers. To determine how prevalent this phenomenon was among different types of neuromuscular diseases, we referenced the Washington University Neuromuscular atlas spinal muscular atrophy image collection^44^ .

Indeed, muscle fibers with similar multiple internalized nuclei were present in samples from bulbospinal SMA, in a Type III SMA patient (age 27), in one SMA low-extremely dominant (LED) patient (age 2), and a TRPV4 mutation SMA case. As the genetic origins of all these diseases is different, it suggests that the multiple internalized nuclei are not a direct consequence of *SMN1* loss but of the resulting muscle pathology.

### No change in global fibrosis or cholesterol levels in SMA Type II muscle

Another common feature of many myopathies is the dysfunction in fibroadipogenic progenitor (FAP) cell population, leading to the accumulation of fibrosis or fat inside the muscle tissue^45^. To assess any potential fibrosis in these tissues we stained the muscle sections with Sirius Red and quantified the intensity of the staining. We observed no overall change between the control and SMA samples (**Figure 2F, Supplemental Figure 2B**).

Metabolic dysfunction and abnormal fatty acid signaling have been previously described in SMA^46,47^, including the accumulation of lipids and cholesterol in some SMA mouse models^47^. In line with these observations, in our cohort, we observed that *LDLR,* the low-density lipoprotein receptor coding gene, was upregulated in SMA samples (**Supplemental Figure 2C)**. This receptor binds to low-density lipoproteins (LDLs) which are the primary carriers of cholesterol in the blood. SMA patients, across all subtypes, are known to have signs of dyslipidemia, including increased blood LDL levels and total cholesterol levels in the plasma and other tissues have been observed in some severe mouse models of SMA ^47^. More recently, abnormal cholesterol accumulation was found in the dystrophic muscles of DMD patients and mouse models^48^ and more studies are uncovering the role of skeletal muscle and fiber type as a global modulator of cholesterol and other lipids ^49^.

Therefore, we decided to assess the total and free cholesterol using the Cholesterol Ester-Glo bioluminescence assay system from the lysates derived from the same muscle samples. However, no differences were observed between total, free or esterified cholesterol (**Supplemental Figure 2D)**. Collectively, this data suggests that the major alterations observed in the SMA Type II muscle are not due to FAP dysfunction, although more detailed analysis is necessary to support this conclusion.

### RNA-sequencing of SMA muscle reveals a heterogenous transcriptional profile post-treatment

To get a broad molecular picture of the state of the treated SMA muscle, we decided to perform RNA-sequencing of each sample. We began by comparing the differentially expressed genes between all SMA patients (n=8) versus all control samples (n=7) (**Figure 3A**), where we observed 154 downregulated genes and 240 upregulated genes (**Supplemental Table 1**). Pathway analysis showed an enrichment for mitochondrial processes such as oxidative phosphorylation and the citric acid cycle (TCA) among the downregulated genes (**Figure 3B**) and an enrichment of calcium signaling and P53 target pathways among the upregulated genes (**Figure 3B**).

**Figure 3:**
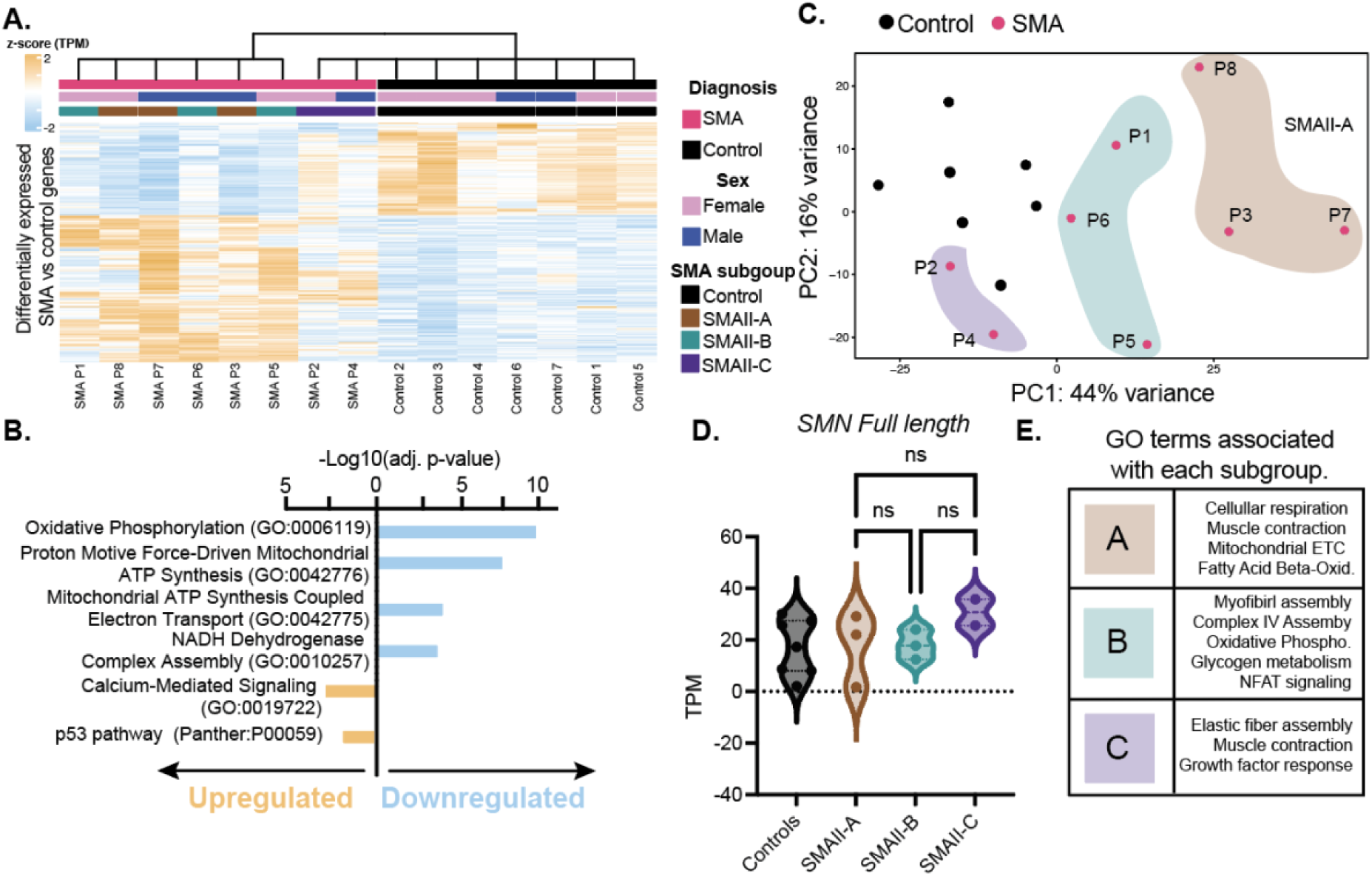
RNA-sequencing characterization of treated SMA muscle cohort. **A.** Heatmap of the genes that are differentially expressed between all SMA muscle samples and all controls. The hierarchical clustering on the top is based only on the differentially expressed genes and shows two groups. By hierarchical clustering two SMA samples, P2 and P4 are included in the control group. Diagnosis, sex and SMA subgroups (based on the top 2000 most variable genes in each sample) are in the first, second and third rows respectively. Transcripts per million are presented as a z-score. **B.** GO terms associated with the upregulated and downregulated genes. **C.** PCA plot obtained using the top 2000 variably expressed genes in each muscle RNA-seq library. The variance explained for each principal component (PC1) is plotted on the axis. **D.** Results from the bulk-RNA sequencing transcripts per million (TPM) for SMN full length transcripts. For controls, the level of *SMN1* was plotted as a reference, while for SMA samples, the levels of *SMN2* were plotted. Means between the three SMA subtypes were compared using one-way ANOVA with multiple comparison testing. **E.** Graphic of the major GO terms associated with the differentially expressed genes in each subcluster of SMA patients.

However, we noted a high degree of variability between different SMA samples (**Figure 3C**). To determine if we could construct molecular subtypes of SMA Type II responses, we used hierarchical clustering on the most variably expressed genes in each sample to construct different subgroups, obtaining three (**Supplemental Figure 3A, Figure 3C**). We named these subgroups SMAII-A (P3, 7 and 8), SMAII-B (P1, 5, 6) and SMAII-C (P2, P4), with SMAII-C having the profile most similar to controls and SMAII-A the least similar (**Figure 3C**). The amount of *SMN2* full-length transcript did not vary significantly between groups (**Figure 3D**). We performed differential expression analysis between controls and each subgroup, obtaining 2722 DEGs in SMAII-A, 475 in SMAII-B and 86 in SMAII-C (**Supplemental Table 1**). Pathway analysis showed an enrichment for mitochondrial oxidative phosphorylation and cellular respiration in SMAII-A and SMAII-B, while SMAII-C showed more enrichment for muscle contractile processes, which was also shared with the other subgroups (**Figure 3E)**.

We next sought to determine if we could observe any sex-effects in our samples, as within the SMA cohort we had an equal number of both sexes (XX, XY, n=4). We also performed this comparison with the control samples. However, only the classic X or Y linked transcripts were differentially expressed, including *XIST, TSIX* up in the female SMA muscle and *TXLNGY* and *DDX3Y* in the male samples (**Supplemental Table 1**).

### Expression of known or hypothesized SMA genetic modifier genes

As is the case for complex diseases, rare monogenic diseases can also be modulated by further genetic factors, giving rise to a lot of variability^50^. Several genetic modifiers have been described for SMA and a handful of these have been validated in human samples^51–53^. Many of these genetic modifiers have been found using genetic screening approaches in *C. elegans*^54^ or *Drosophila* ^55^. To determine the effect that these modifier genes might have on the muscle, we assembled a list of known and hypothesized SMA modifiers (**Supplemental Table 1**) and assessed their expression in our samples (**Supplemental Figure 3B**). Hierarchical clustering of the samples revealed that all but the two samples from group SMAII-C (P2 and P4), clustered away from the control samples (**Supplemental Figure 3B**). However, two other groups, SMAII-A and SMAII-B were not characterized by a specific modifier gene expression, suggesting that the influence of these genes were not driving the overall effect in the transcription. Of particular interest, we observed an increase in NCALD expression in SMAII-A samples. This trend is consistent with previous studies reporting that a decrease in NCALD expression was associated with milder SMA phenotype ^56^. We saw a similar elevation in *NAIP* levels in SMAII-A samples, although the importance of this modifier gene is not well understood. Finally, we saw no change in PLS3 expression between the different groups ^57^ (**Supplemental Figure 3C)**.

### Mitochondrial oxidative phosphorylation complexes are altered to varying degrees in Type II SMA Muscle

Mitochondrial metabolic capacity is an important key to muscle function. We thus decided to more closely investigate the differential expression of mitochondrial genes between the three patient subgroups. We compared the differentially expressed genes in each group (**Supplemental Tables 1**) to the Mitocarta3.0^58^. Of the 1136 genes listed in the MitoCarta3.0, 42% of the genes were differentially expressed in the severe SMA, while only 6% and 0.5% of them were differentially regulated in the SMA II-B and SMAII-C, respectively, showing the stark difference in mitochondrial involvement between the subtypes (**Figure 4A**). In the SMA Type II-A subgroup, we observed a broad downregulation of Complex I-V subunits (**Figure 4B**) and predicted metabolite change analysis showed impingement of NADH levels (**Figure 4C**). Decreases in mitochondrial function, density and expression of electron transport change (ETC) genes have already been reported in several SMA models, including in the spinal cord and muscle tissue to Type I patients^59,60^. Indeed, comparing out results to a previously published muscle-only *Smn1* knockout mouse model^61^, we observed a modest overlap between the genes downregulated in the SMAII-A group and in the mouse muscle (13% of downregulated genes in SMA II-A). Predominantly, these pathways were associated with mitochondrial matrix proteins and other metabolic processes (**Supplemental Figure 3F, Supplemental Table 1**).

**Figure 4:**
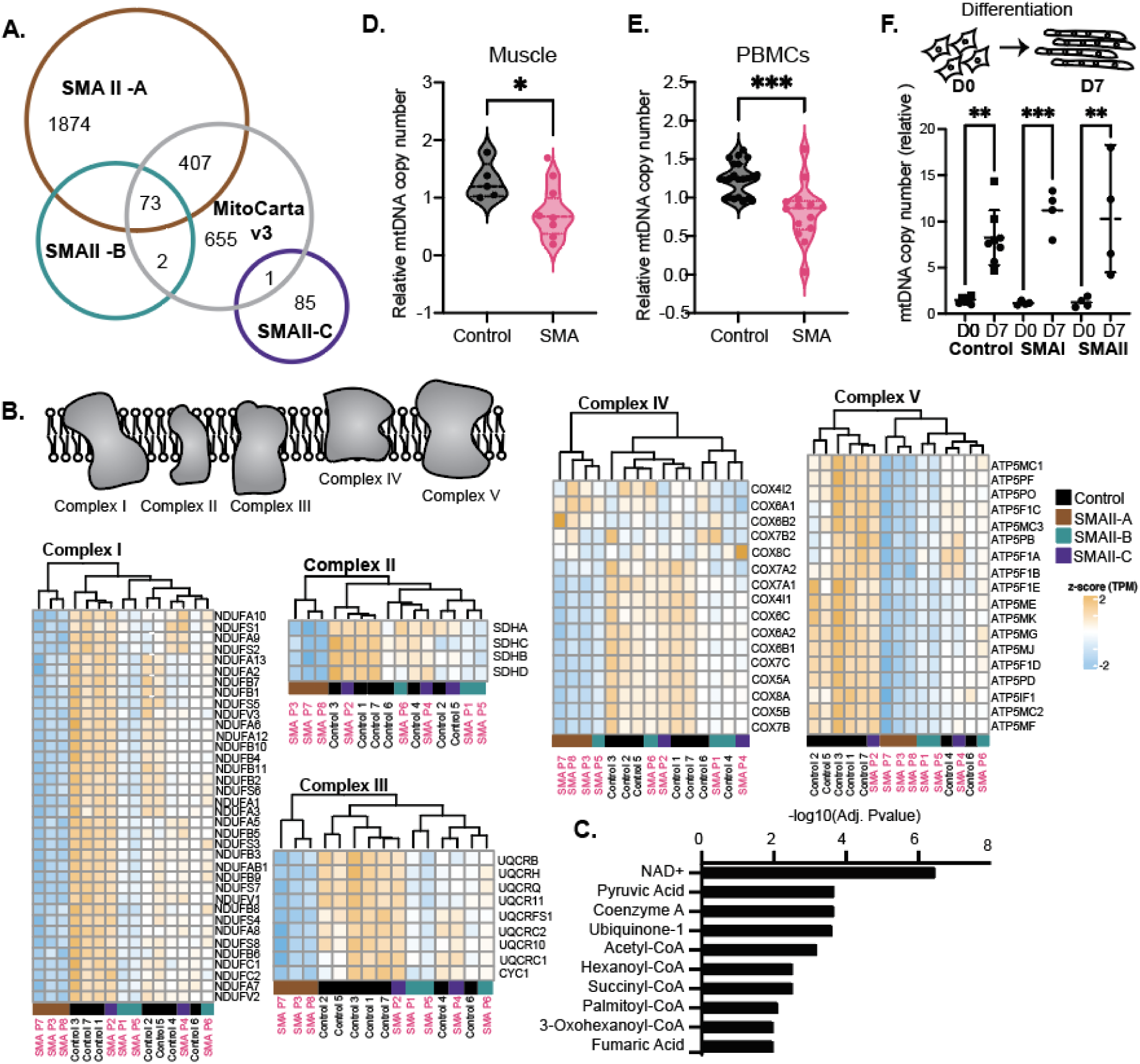
Mitochondrial DNA copy number is lower in Type II SMA muscle but is not an intrinsic feature of SMA deficiency in immortalized myoblasts. **A.** Overlap between the genes known to be localized in the mitochondria, from MitoCarta v3 compared to the differentially expressed genes in each subgroup of SMA Type II muscle. **B.** Scheme of each of the complexes of the ETC, with the corresponding genes diagrammed in the heatmaps below. Hierarchical clustering was performed based on the expression of each subset of complex genes. **C.** Predicted metabolite alterations based on the differentially expressed gene profile of the subtype SMA II-A. **D.** Relative mitochondrial copy number, normalized to the numbers of genomic DNA amplified gene, B2-microglobulin in paravertebral muscle. Each point represents one sample. Groups were compared with the student’s t-test. * <0.01 **E.** Relative mitochondrial copy number, as above, in DNA from PBMCs. Each point represents one sample. Groups were compared with the student’s t-test. ** <0.001. **F.** (Top). Scheme of myoblast to myotube differentiation. (Bottom). Mitochondrial copy number as above in DNA derived from myoblasts at Day 0 and Day 7 of differentiation. Groups were compared using a one-way ANOVA with multiple group comparisons. ** <0.001, ***<0.0001.

Given the notable changes we observed in mitochondrial processes, we first hypothesized that this might reflect a change in the fiber types, as mitochondrial state and dynamics are key features of the two different types of muscle fibers^62^. Thus, we decided to determine the fiber type composition of our muscle samples. It has been previously demonstrated that muscle samples from patients undergoing spinal surgery accounted for 74% of Type I (slow) fibers in the superficial and deep thoracic regions and that in the lumbar region, 57% of the fibers were type I in the superficial muscles and 63% were Type I in the deep muscles^63^. We utilized a bulk RNA-sequencing deconvolution method, where bulk RNA-seq data can be used to predict the fiber type in the original tissue sample, based on the expression profiles of Type I (slow) and Type II (fast) muscle fibers derived from single-cell sequencing profiles^64^. In accordance with these previous findings, fiber type deconvolution showed that in control samples, we had an average of 66% Type I fibers (**Supplemental Figure 3C**). On the other hand, in SMA samples we had an average of 69% Type I, suggesting that overall fiber type was not highly affected in the paravertebral muscle we sampled (**Supplemental Figure 3C**). Previous studies on SMA patients have found a loss of Type II glycolytic fibers^65^, which mirrors what is seen in mouse models of SMA ^11,66,67^. However, in all these studies, muscle with a higher starting fraction of Type II fibers were investigated. We plotted the expression of different myosin heavy chain forms across different patients (**Supplemental Figure 3D**), and overall, we observed no clear trends in myosin heavy chain changes between the control and SMA groups.

### Mitochondrial DNA copy number is lower in treated SMA samples than in controls

We next sought to test the number of mitochondrial DNA (mtDNA) copies in the SMA muscle samples. mtDNA copy number is used as a surrogate measure for the number of mitochondria, when such measurements are not possible. Using a previously established protocol for a qPCR quantification of mitochondrial DNA copy number^28^, we measured mtDNA copy number from the extracted DNA of the same samples used for sequencing. We observed that globally mtDNA was significantly lower in SMA than in control samples (**Figure 4D**), although this had no correlation with SMA group. To test if this was a phenomenon that was more broadly true in SMA pathophysiology, we performed the same measure on DNA extracted from the circulating PBMCs of Type III patients, all of whom had been treated with Nusinersen and Ridisplam, and likewise noted a significant decrease in mtDNA copy number (**Figure 4E**). Collectively, these results suggest that mitochondrial loss is a property of SMA tissues.

This led us to question if mitochondrial regulation was a byproduct of the loss of SMN. To test this, we utilized immortalized myoblasts derived from SMA patients (Type I and Type II) or healthy controls. However, we did not see a difference in mtDNA copy number, despite a difference in *SMN* expression (**Supplemental Figure 3F**). This is true both among the SMA myoblasts, but also among different control cell lines that had different expression levels of SMN1/3 (**Supplemental Figure 3F**). To test if this discrepancy with the *in vivo* muscle samples was due to the immaturity of the myoblasts, we performed a serum-free differentiation for seven days to generate myotubes. It was been previously described that myoblasts increase their mitochondrial DNA copy number and number of mitochondria as part of the differentiation processes^68^, and we could recapitulate this finding in our control myoblasts (**Figure 4F**). The SMA myoblasts were also able to increase their mtDNA copy number to the same degree as the controls (**Figure 4F**). The results mirror what was previously seen in myoblasts differentiated from SMA ESC lines, where copy number did not change, although ATP production capacity was altered^69^. Taken together, this data suggests that mtDNA copy number changes are not an intrinsic feature of SMN loss, but rather a downstream consequence of developmental delays or alterations that occurred in the early SMN-less environment before treatment and that may be leading to the increased atrophy and altered myofiber architecture that we observe.

### P53 target gene activation is differential between different SMA subtypes

DNA damage response and cellular stress are also major hallmarks of SMA pathology ^70–72^. Thus, we sought to investigate if SMN-targeted therapies could ameliorate these features. One of the pathways activated when comparing all SMA samples versus all controls (**Figure 3B**) is P53 signaling. Importantly, this implies that the damage response observed in these SMA samples goes beyond any damage response associated with the scoliosis, as the control samples were non-SMA but were still undergoing spinal surgery. In all samples we observed an increase in P53 targets: *GADD45A, GADD45B, SERPINE1, TNFRSF10B, PPM1D and CCNG1* (**Supplemental Figure 4A**). Although *TP53* was not differentially expressed when we compared all SMA samples to all controls, we could see an increase in *TP53* in the SMAII-A, those samples which are most different from the controls (**Supplemental Figure 4B**).

P53 has been found to be activated in several SMA models and has been associated with the MN death that characterizes SMA ^70,71,73^. To better understand any potential p53 dependent regulation in our muscle samples, we first downloaded the known transcriptional targets from the *TP53 Database*^74^. From this database, we assembled a list of 343 validated p53 target genes. These targets are taken from a wide variety of tissues, thus their specificity to muscle is not validated. We overlapped these genes with the differentially expressed genes in the SMA samples and found 17 genes (5% of all database targets) that overlapped (**Supplemental Figure 4B**). However, due to high degree of variability, we also compared the overlap by group. In the SMAII-A group, we found that 66 P53 target genes (20% of the target list) were affected, while in SMAII-B, we observed 11 genes that overlapped, all shared with SMAII-A, and in SMAII-C there was only one target gene altered (**Supplemental Figure 4C**). Determine if P53 was activated as a response to DNA damage, we performed a western blot for ser15 phosphorylation on P53. Serine 15 is the primary target of the DNA damage response and can be phosphorylated by ATM and ATR protein kinases^75^. However, although some samples appeared to have particularly elevated levels, P53 activation was not globally higher, and did not correlate with the previously established transcriptional profiles (**Figure 3C**). More studies will need to be performed on P53 activation to determine if it is a major player in post-treatment SMA muscle pathology.

### Mitochondrial regulatory microRNA mirR-1 but not mir-206 is misregulated in post-treatment SMA Type II PV muscle

Given that our fiber type analysis suggested that selective loss of different fiber types was not responsible for the differences in the mitochondrial pathways that we observed, we next explored other known mechanisms of mitochondrial regulation. These can include changes in mitochondrial fusion and fission dynamics, changes in mitochondrial DNA biogenesis, mitophagy or proteostasis or changes to mitochondrial DNA replication (**Figure 5A**). However, analyzing our differentially expressed genes, both in all the SMA samples and by subgroup, we did not observe gene expression changes that would reflect changes in these pathways (**Supplemental Table 1**).

**Figure 5:**
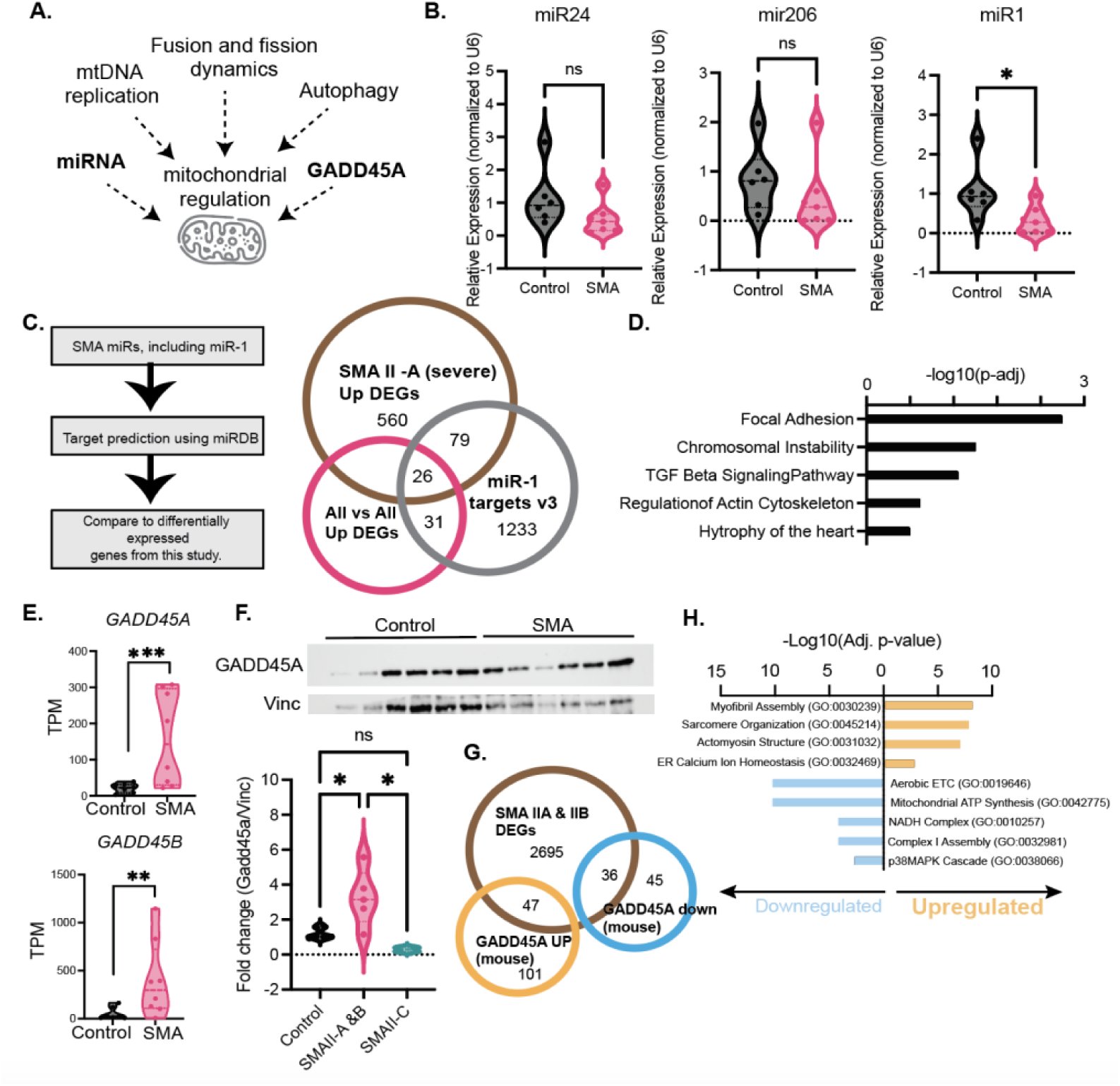
Misregulation of miR-1 and GADD45A in Treated PV SMA Muscle. **A.** Scheme of previously proposed mechanisms of mitochondrial regulation. **B.** Relative quantification of miR24, miR206, and miR1 respectively. Each point represents a muscle sample. Expression was normalized to the expression of the U6 snRNA. Comparisons between the two groups we performed using a student’s t-test. * <0.01. **C.** (left) Scheme of the process used to identify which genes, if any, in the differentially expressed genes were associated with known changes in miRNAs in SMA. (right) Overlap between the predicted targets of miR-1 using miRDB with the genes that are differentially expressed in all the SMA patients or in the SMA II-A group. **D.** Pathway analysis associated with the genes that are differentially expressed in the SMA samples that are also predicted targets of miR-1. **E.** Transcript per million (TPM) counts for *GADD45A* and *GADD45B* from the RNA-sequencing of muscle samples. Each point represents one patient sample. P-values are based on the adjusted p-values derived from DeSEQ2 which considers multiple hypothesis testing. **F.** Representative western blot for GADD45A SMA and control muscle lysate compared to a vinculin housekeeping control. Bands are quantified on the right. Sample groups are compared using a one-way ANOVA with multiple comparison testing. **G.** Overlap of the genes that are differentially expressed in the SMA II-A and II-B groups with the empirically determined GADD45A target genes in the mouse muscle^72^. **H.** Pathway analysis for the GADD45A target genes determined from the mouse muscle that overlap with differentially expressed genes in the SMA II-A and II-B samples. n.s.: P . 0.01, * <0.01, ** <0.001, ***<0.0001.

MicroRNAs (miRNAs) have also been shown to have an important role in mitochondrial function, especially in the regulation of transcription and translation. Indeed, many miRNAs, such miR-1, can be directly imported into the mitochondria^76^ and many have well described roles in muscle development, and are aptly named myoMiRs ^77–79^. Perhaps the best established of these is the role of miR-206 in skeletal muscle maturation and specifically mitochondrial health^80,81^. Furthermore, many studies in SMA have highlighted the essential role of alterations of miRNAs to the SMA phenotype, and these miRNA have also been proposed as biomarkers of Nusinersen response and disease progression ^78,82–87^. If changes in miRNA expression observed in SMA are a direct or indirect consequence remains to be determined, although one study suggests that SMN is involved in the direct transcription of some microRNAs^82^.

We reasoned that given our restored SMN levels, we could determine if these prominent myomiRs were altered in our samples. We used microRNA amplification and probe sets to measure the levels of three important myomiRs which have been implicated in SMA: miR1, miR26 and miR206. No difference was observed in the levels of miR-24 and miR-206. However, the levels of miR-1 were significantly decreased in the SMA samples, regardless of the previously established subtype (**Figure 5B**). This is in-line with previous reports that metabolic maturation in muscle stem cells in partially achieved by miR-1 mediated inhibition of the Dlk-1-Dio3 locus, which contains miRs that repress mitochondrial gene expression^88^. One recent study found that SMN, in combination with MYOD, can regulate the expression of miRNAs that are related to mitochondrial gene regulation, including miR-1^82^. However, we observed this decrease despite detecting similar levels of SMN protein (**Figure 1 C**), suggesting additional levels of regulation.

To better understand the effect of the differential miR-1 regulation we observed in our muscle samples, we used the same miRDB database to predict it’s targets, both for has-mir1-5p and has-mir1-3p as our primers cannot distinguish between these two forms. This resulted in a theoretical 1233 target genes (**Figure 5C**). We then compared these to our list of all differentially upregulated genes (240 genes) and found that of these 15% (38/240) were predicted miR-1 target genes. Comparing the more severe SMA cases in group SMAII-A, which had 804 upregulated genes, we found an additional 79 targets, plus 26 targets that were shared with those previously found (**Figure 5C**). Using the list of 104 target genes in the SMAII-A group, we performed pathway analysis and found target genes in pathways related to focal adhesion, chromosomal instability, TGF-beta signaling, and hypertrophy (**Figure 5D**). This further validated by previous reports on the role of miR-1 in TGFB signaling and cardiac hypertrophy ^89^.

### GADD45A RNA and protein levels are elevated in treated SMA type II muscle

Increased levels of GADD45A are a well described feature of muscle atrophy^90–92^. One recent report linked GADD45A activation to decreased mitochondrial function (**Figure 5A**)^92^, and as we could observe transcriptional activation of *Gadd45a* and *Gadd45b* across our SMA samples (**Figure 5E**), we hypothesize that this might be linked to the mitochondrial deficiency we observed. We validated the increase in GADD45A protein using western blots. GADD45A levels were generally elevated in samples from SMAII-A and SMAII-B but not from group SMAII-C (**Figure 5F**) suggesting that the GADD45A activation scales with the amount of mitochondrial oxidative phosphorylation loss we observed.

To determine if GADD45A elevation is a feature of SMA, we used our severe-to-intermediate mouse model of SMA, the *SMNΔ7* model. These mice have disease onset between postnatal day 5-7 (P5-7), with an average lifespan between 14-16 days (P14-16). We observed that at both P7, the onset of motor symptoms, and P14, the end-stage of the disease, there was no significant change in GADD45A levels in the SMND7 muscle. This led us to conclude that GADD45A activation is not an intrinsic feature of SMA, but a downstream consequence of long-term muscle development in an SMN-low environment.

To determine if GADD45A activation, which is known to play a role in transcriptional regulation in response to atrophy, was involved in the gene expression changes we observed in the SMA muscle samples, we compared the differentially expressed proteins induced after GADD45A overexpression in the mouse *tibialis anterior* muscle ^92^ to the differentially expressed genes in SMA II-A and SMA II-B, as SMA II-C samples appeared to have little to no levels of GADD45A induction (**Figure 5F, Supplemental Figure 4A**). We observed that 44% of the downregulated GADD45A targets and 32% of the upregulated GADD45A targets were also differentially expressed in the SMA muscle (**Figure 5F**). Pathway analysis showed that the downregulated genes included oxidative phosphorylation and other mitochondrial metabolism processes, while the upregulated genes related to myofiber and sarcomere organization and actin polymerization (**Figure 5F**). This suggests that GADD45A may have both protective and detrimental effects on the SMA muscle.

## Discussion

Here we present a molecular characterization of a cohort of eight SMA Type II muscle tissue samples, collected from adolescent aged patients after treatment with SMN-targeted therapies. We observe that SMN protein and RNA levels in muscle are restored after treatment with Nusinersen and Ridisplam to levels at or above that of control samples. Despite this, we detect molecular and phenotypic features of the muscle that are not restored. Notably, this is characterized by fibers with multiple internalized nuclei, a decreased mitochondrial copy number count, and a transcriptional profile consistent with a reduction in oxidative phosphorylation capacity. Additionally, we see a persistent activation of GADD45A in a subset of samples, a mediator of muscle atrophy and mitochondrial loss in muscle. It is also important to note that the SMA samples are compared to controls that are also undergoing scoliosis surgery, allowing us to find define hallmarks of SMA muscle beyond any changes associated with scoliosis^93^. Collectively, this work suggests that to fully restore SMA muscle functionality, targeted combination treatments need to be designed in addition to SMN-based therapies that are currently available. These may be considered *SMN-irreversible* processes^94^, although their direct link to SMN loss during development remains to be understood.

### Reactivation of SMN levels

State-of-the-art SMA therapies directly seek to increase the amounts of SMN protein. Both Nusinersen and Risdiplam are SMN dependent therapies whose mechanism of action is based on inducing the inclusion of exon 7 in the remaining *SMN2* copies^1,14,15^. However, the effectiveness of these therapies in peripheral tissues remains poorly understood. A previous study on *SMN* mRNA splicing in several tissues following ASO injection had found induction of proper splicing only in the spinal cord but not in the CNS or other peripheral tissues of treated SMA patients, including the iliopsoas and diaphragm muscles ^24^. However, that study was based on post-mortem samples from young children (<72 months), with two copies of SMN2, consistent with an SMA Type I profile ^2,24^. Many of these samples were procured from very severe cases not long after the first injection. By contrast, although our study focuses on a different group of SMA patients, we can see a correction of the splicing defect, although we did not validate the penetration of the antisense oligonucleotide directly. Our data suggest that at least in peripheral tissues that are close to the injection site, i.e. paravertebral muscle, the SMN-centric therapies can act as desired. Additionally, it suggests that the other phenotypic changes we observe may require combination therapies with non-SMN modes of action.

### Heterogeneity among SMA muscle samples

Our study has the limitations of an observational human study, and particularly, as in many rare-disease studies, there are a limited number of patient samples. To add to the complexity of the small sample size, the type of treatment and frequency, and number of years under treatment are different with each patient and the full natural and clinical histories are not available for all samples. This increases the complexity of interpreting the data, as not all variables can be effectively accounted for. Despite these challenges, we were able to collect high quality data, with minimal variation among the control samples, and to capture the expected percentage of Type I vs Type II fibers, suggesting that any observed heterogeneity is not due to sampling bias.

We were able to construct a transcriptional signature of SMA Type II muscle after treatment, with no differences observed based on sex. Comparing the control samples to Type II SMA samples, we observe a higher degree of variability in the SMA samples, suggesting that to some degree this heterogeneity is related to the evolution of the pathology. We could identify three subgroups of SMA patients, II-A: those with the most severe loss of ETC complex, especially that of complexes I-III, II-B: a subset with intermediate mitochondrial protein involvement and changes to cytoskeletal proteins and subset II-C which clusters generally with controls, with some alterations in cytoskeletal proteins. However, both patients in the II-C group had multiple internalized nuclei, suggesting that the altered pathways have functional consequences. We could also observe an activation of the *P53* pathway and GADD45A expression in a group dependent manner, with higher degree of activation observed in the more severe II-A and II-B groups. Although further studies are needed to validate these findings, it suggests that one subset of SMA patients could benefit from mitochondrially targeted therapies while the other from cytoskeletal targeted therapies.

One possible source of the subtype variability could be genetic modifier genes, as several have been well described to influence SMA presentation. We observed an upregulation of *NCALD* in the SMA cohort, with the notable exception of the two II-C patients, where it remains around control levels. This is consistent with previous reports that decreasing *NCALD* expression ameliorates SMA phenotypes ^56^. The expression of the other well described modifier, *PLS3* ^57^, was also highly variable, with the vast majority of SMA samples not showing an upregulation above the variation observed in the control cohort. Further sources of epigenetic variation remain to be explored.

### Multiple internalized nuclei characterize a subset of SMA samples

Another striking feature we observed in the SMA muscle histology was the presence of multiple centralized nuclei, present in 3 out of 5 patients we were able to perform histology on. The fusion, alignment and movement of myonuclei to the periphery is a highly coordinate process, in which many players remain poorly characterized ^95,96^. In 3/5, we observed multiple internalized, though not perfectly central, nuclei, particularly in fibers with a large diameter. Based on these observations, one speculation may be that in these fibers, satellite cells fusion is an active process, perhaps in response to damage signal, but once fused the myonuclei are not able to move to the periphery because of the structural defects on cytoskeletal components, resulting in massive fiber diameters. The peripheral position of nuclei along the muscle fiber is a hallmark of skeletal muscle tissue. Actin dynamics have also been implicated in nuclear movement to the periphery, and it is known that actin assembly can organize the desmin network necessary for myofibril crosslinking^97^. Differential gene expression analysis showed several actin related pathways in SMA patients with multiple internalized nuclei. However, we speculate that the mechanism responsible for these observed multinucleated fibers is not the same as in centronuclear myopathies caused by mutations in *BIN1*, MTM1 or DNM2, as classically those result in the persistence of myofibers with a single, well-centralized nucleus^98,99^. Further characterization of affected fibers using special transcriptomics approaches may elucidate putative mechanisms. If this effect is dependent on *SMN1* loss or not remains to be determined, however, to date, we are unaware of other reports of multiple internalized nuclei in SMA muscle histology, including in our own studies, although large, hypertrophic fibers have been seen in other histological analyses^100^.

### Involvement of *SMN1* in mitochondrial homeostasis

Mitochondria are essential organelles present in all eukaryotic cells that provides most of the ATP production via oxidative phosphorylation. This process, which occurs along the inner mitochondrial membrane, is mediated by the electron transport chain (ETC), a series of protein complexes that turn chemical potential gradients into chemical bonds. Additionally, mitochondria serve as Ca2+ storage, can be a source of reactive oxygen species (ROS) and are mediators of many cell stress and apoptosis responses. Due to their critical requirement of oxygen, and their large consumption of cellular oxygen, they are also critically involved in the sensing of oxygen levels and hypoxia. Likewise, hypoxia is major feature of Type I SMA^101,102^. The precise role of *SMN1* in mitochondrial health and homeostasis remains unclear ^103^. There is controversial evidence that *SMN1* may transiently localize to either the inner or out mitochondrial membrane, as well as proposed roles for SMN in the regulation of mitochondrial shape or trafficking, via actin regulation, or through mitochondrial gene expression regulation via miRNA production.

For example, in a muscle-specific knockout of *Smn1*, the accumulation of mitochondria with morphological abnormalities and with impaired complex I and complex IV activity and respiration deficits was observed^61^. Similar results were obtained from SMA motor neurons^104,105^, where the authors also noted that these complex I deficiencies can cause the accumulation of ROS, which has cascading effects on other protein homeostatic processes ^105^. In the in vitro model system C2C12, SMN has been proposed to target, in combination with MYOD, two key miRNAs for mitochondrial health: miR-1 and miR-206 ^82^. Similar alterations in the muscle’s metabolic processes, measured by MRI after muscle stress tests, were obtained in Type III and IV SMA patients ^106^. In particular, these MRI images showed that in the triceps, there was a disproportionate loss of the fastest contracting myofibers and decreased mitochondrial ATP synthetic function in residual white myofibers ^106^. Here we find clear evidence that among a subset patient, there is a downregulation of most of the complexes of the ETC (SMA Type II-A) and that in the second subset, we see downregulation of these same complexes, but not to the same extent.

Finally, among the last group, there is not clear downregulation. Finally, we observe lower mitochondrial DNA copy numbers, often used a proxy for mitochondrial numbers in both muscle and PBMCs from treated SMA Type II and Type III patients, respectively. However, in myoblasts derived from Type I and Type II SMA patients, we do not observe a lower mitochondrial number, suggesting that the phenomenon is not directly related to *SMN1* loss, at least not *in vitro*. Additional factors – be they genetic or epigenetic – maybe determine the degree of mitochondrial involvement, or this mitochondrial loss may be a result of development in an *SMN* -low environment.

### Conclusion

Taken all together, the findings of this study support the hypothesis there are fundamental developmental delays that occur in Type II SMA patients’ muscle which develops in a low SMN state, prior to the onset of therapy. Our data lends strength to other clinical findings that early intervention and treatment, even before the onset of symptoms in Type II and III patients is crucial, as these early developmental delays do not appear to be rectified by normal SMN levels postnatal. Additionally, it argues for the need for compound therapies that target these problems, regardless of their SMN independent nature. Finally, it brings up the biological and clinically interesting questions of why Type IV adult patients have symptoms only later in life and how the roles of SMN with aging differ from those of SMN in development.

## Data availability

Raw and processed RNA-sequencing files are available in GEO under accession number GSE252128.

## Author contributions

FCG – designed the research study, performed experiments, analyzed, and interpreted data, generated figures, and wrote the manuscript.

S.A. – performed experiments.

S. P. – performed experiments.

E.G. – performed experiments.

S.M. – performed experiments.

M.M and S.V. collected and preserved the muscle samples.

K.M. - derived the immortalized cells used in this study.

P.S – designed the research study, oversaw data acquisition and interpretation, and wrote the manuscript.

## Acknowledgements

We thank the patients and their families for consenting to the donation of tissue samples to this research. We would also like to thank the members of the Smeriglio lab for their helpful discussions and the platforms of MyoBank and MyoLine which make the collection and generation of patient derived samples possible, as well as MyoImage, which supports the microscopy center. We would like to thank the Genethon DNA Bank for the DNA from Type III treated SMA patients. We would like to thank the staff of IGenSeq Platform (Institut du Cerveau) for their help with the RNA-sequencing library quality control and use of their facilities.

---

**Supplemental Figure 1:**
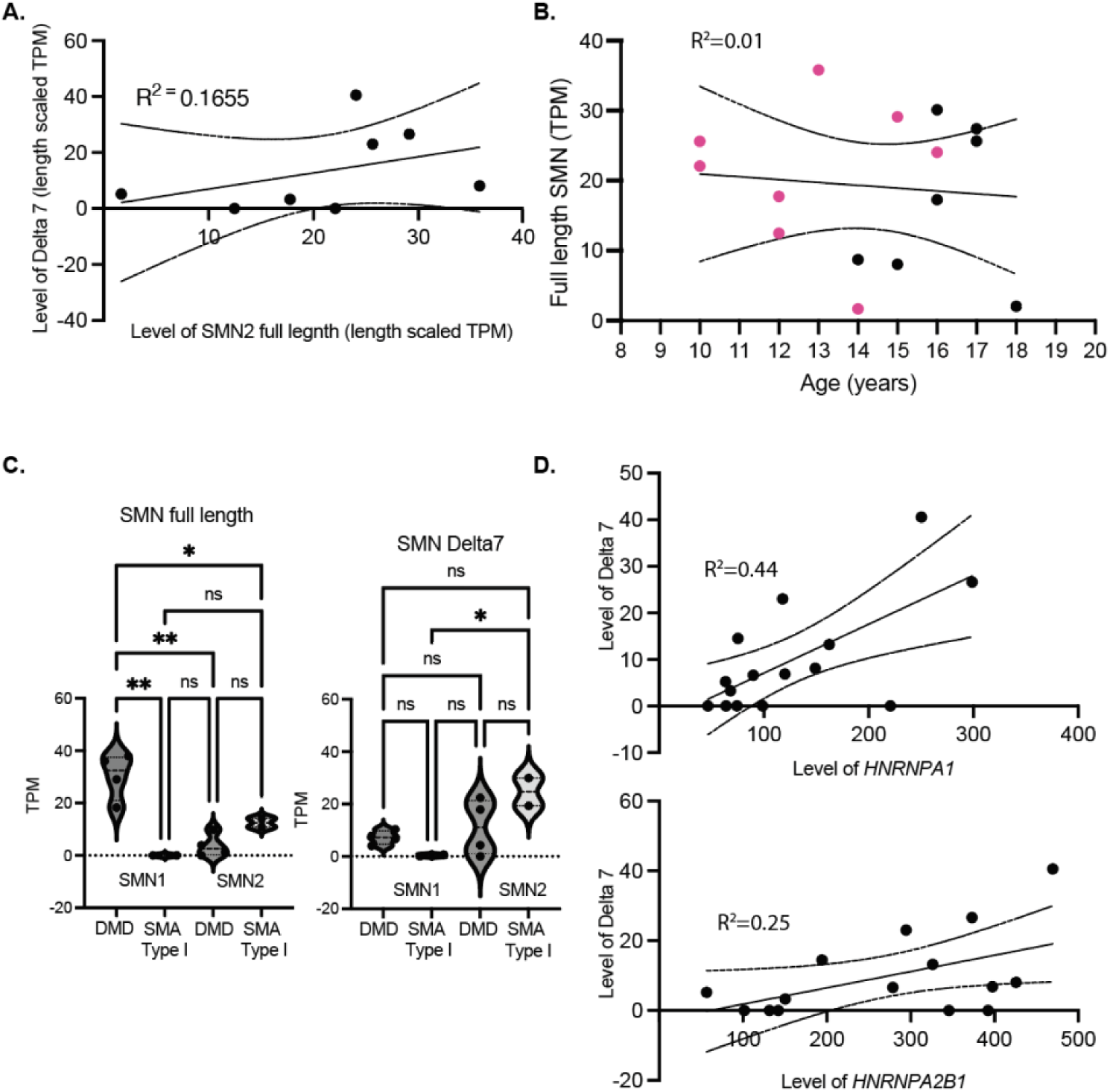
Regulation of SMN expression. **A.** Relationship between *SMN2* transcripts per million (TPM) and Delta7 transcripts from both loci. There is no significant relationship between the two as assessed by a linear regression. 95% confidence intervals for the linear analysis are shown and R-squared goodness of fit values are shown. Each dot represents a single muscle sample. **B.** Relationship between age and total amount of full-length SMN transcript from both loci. There is no significant relationship between the two as assessed by a linear regression. 95% confidence intervals for the linear analysis are shown and R-squared goodness of fit values are shown. Each dot represents a single muscle sample; SMA samples are designated in pink and control samples are in black. **C.** Reads, represented as transcripts per million (TPM) mapping to the SMN full length transcript (left) or the SMN delta 7 transcript missing exon 7 (right). The origin of full-length transcript, either the *SMN1* or *SMN2* locus, is designated below each pair of violin plots. Data was reanalyzed from GSE97806. Each point represents a sample. P-values derived from DeSEQ2 differentially expression analysis and represent adjusted p-values for multiple hypothesis correction. * <0.01 ** <0.001. **D.** TPMs for the delta 7 transcript in control and SMA muscle samples, compared to the TPMs of HNRNPA2A1 (top) and HNRNPA2A1 (bottom). 95% confidence intervals for the linear analysis are shown and R-squared goodness of fit values are shown.

**Supplemental Figure 2:**
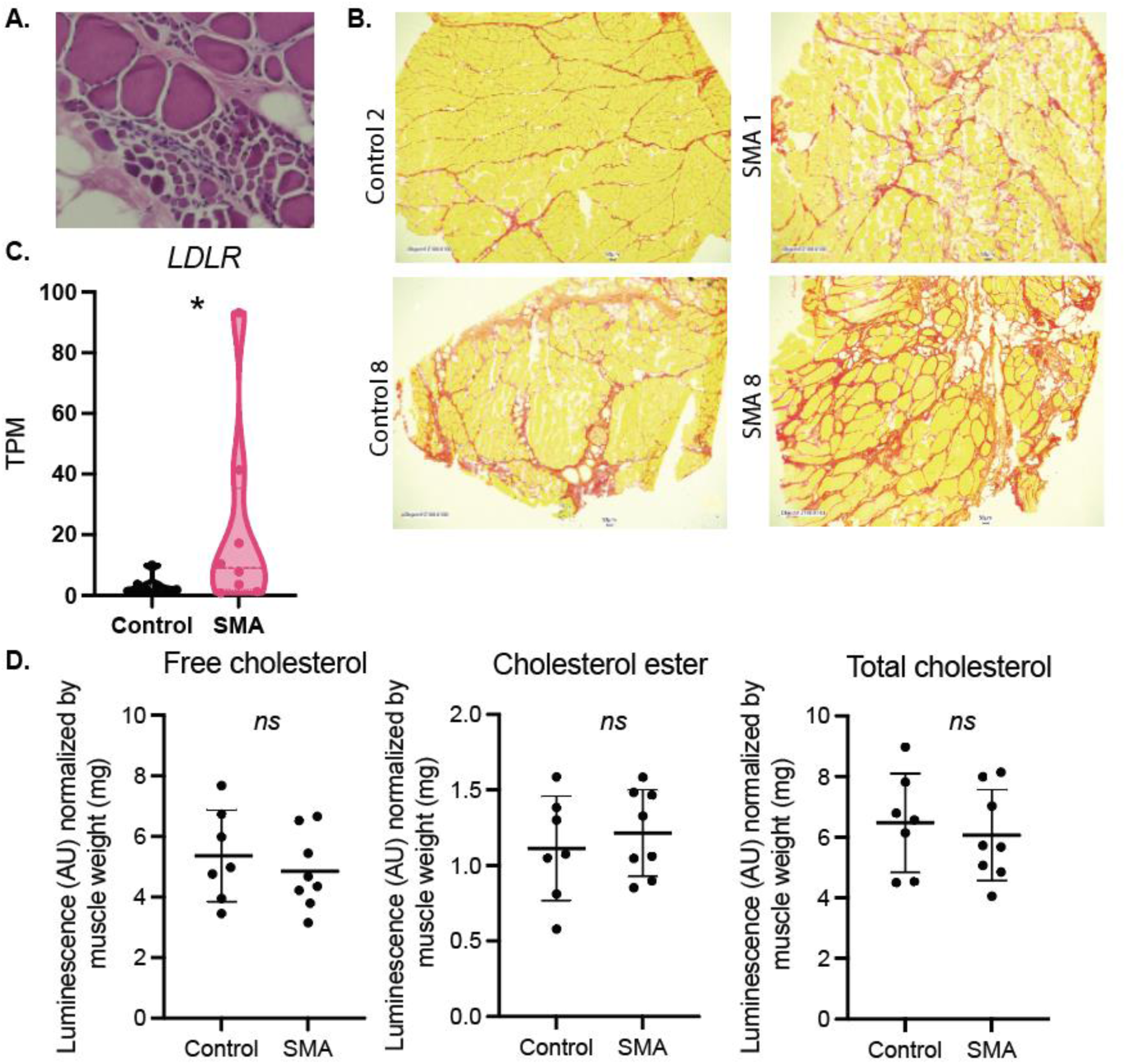
Cholesterol and fibrosis are not increased in treated SMA Type II Paravertebral Muscle. **A.** Zoom in from SMA Patient #2 (reproduced from Figure A) to highlight the clusters of small fibers. Of the patient samples tested, this was the only sample that had this pattern. **B.** Representative Sirius Red staining images on control and SMA muscle samples, quantified in Figure 4. **C.** Transcript per million (TPM) counts for seven LDLR from the RNA-sequencing of muscle samples. Each point represents one patient sample. P-values are based on the adjusted p-values derived from DeSEQ2 which considers multiple hypothesis testing. **D.** Cholesterol-Glo luciferase assay. Each point represents one muscle lysate. Groups were compared using students-test test. *: p <0.01.

**Supplemental Figure 3:**
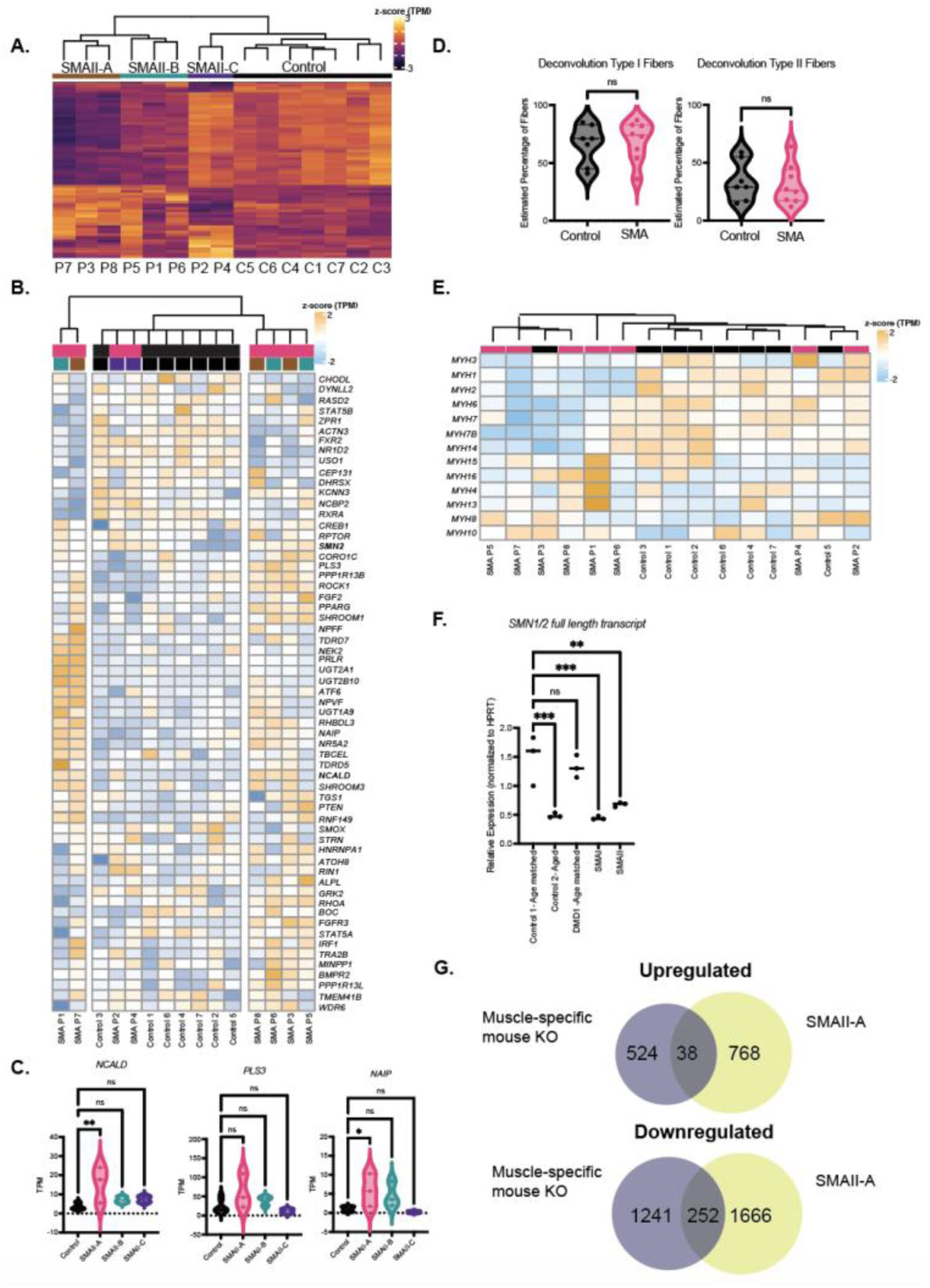
Transcriptional Heterogeneity in Type II SMA Paravertebral Treatment after treatment. **A.** Heatmap showing the genes that are differentially expressed in each SMA subgroup. Hierarchical clustering is performed using the differentially expressed genes. **B.** Heatmap of the expression of putative and validated SMA modifier genes. Hierarchical clustering is based only on gene expression. Transcripts per million are presented as a z-score. **C.** Transcripts per million (TPM) of three well-known SMA modifier genes, *NCALD, PLS3* and *NAIP.* TPMs were compared using a one-way ANOVA with multiple comparison testing comparing all samples to the controls. Each point represents a single muscle sample. P values = * <0.01, ** <0.001, ***<0.0001. **D.** Results from the bulk-RNA sequencing fiber type deconvolution for Type I and Type II fiber signatures. Comparison was done with a student’s t-test. Each dot represents one sample’s RNA-seq library. **E.** Heatmap of the expression of myosin heavy chains in each sample. Hierarchical clustering is based only on myosin expression. Transcripts per million are presented as a z-score. **F.** Real time quantitative PCR of SMN1/2 transcript expression in myoblasts. SMA cells were compared to both age-matched control (10-16 years old) and DMD1 sample, as well as to an older non-aged matched control (50-63 years old). **G.** Comparison between the differentially expressed genes in a muscle-specific *Smn1* knockout mouse (KO) compared to the genes differentially expressed in the SMA II-A subgroup.

**Supplemental Figure 4:**
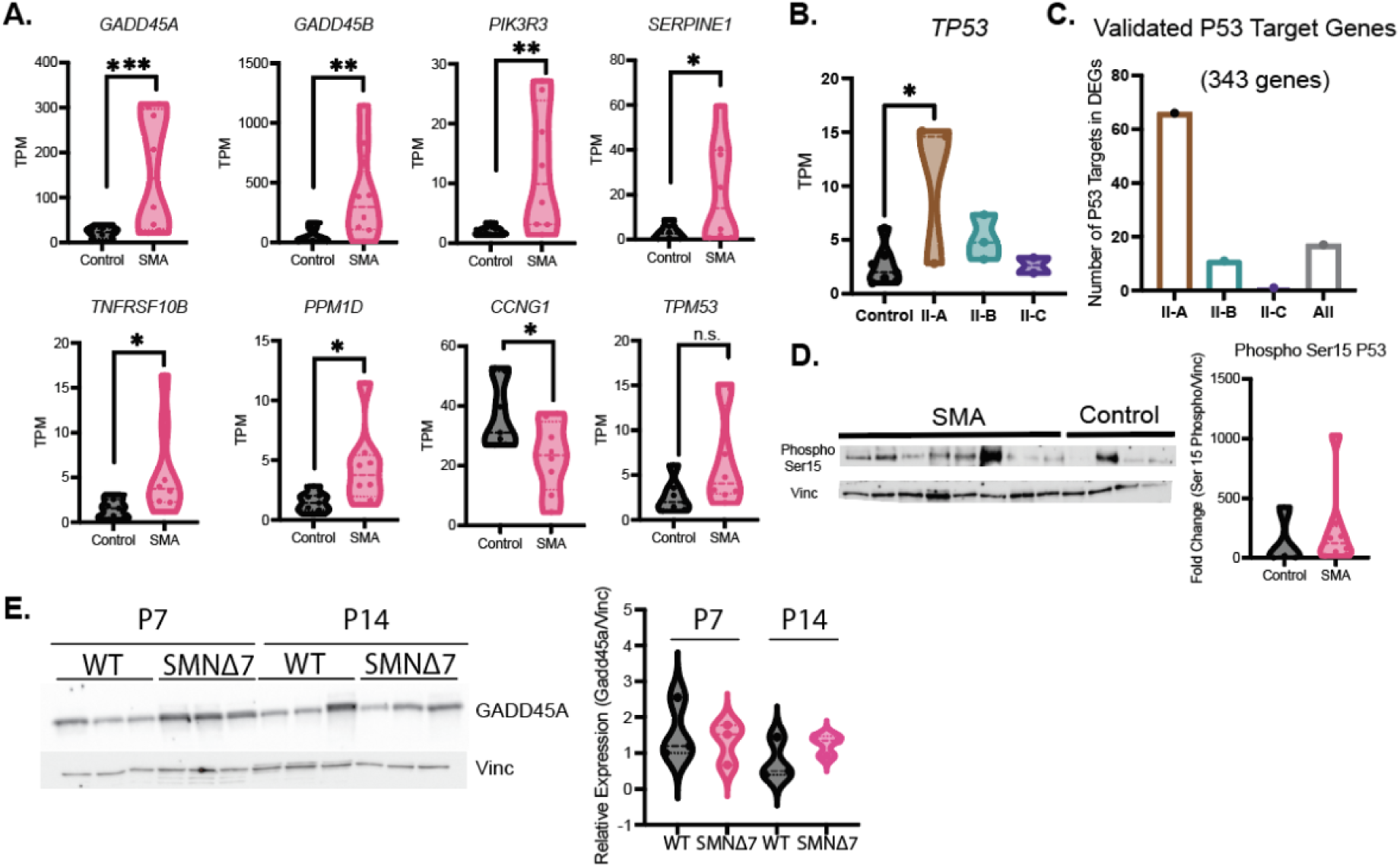
P53 Targets in SMA Muscle. **A.** Transcript per million (TPM) counts for seven P53 target genes and *TP53* from the RNA-sequencing of muscle samples. Each point represents one patient sample. P-values are based on the adjusted p-values derived from DeSEQ2 which considers multiple hypothesis testing. *GADD45A* and *GADD45B* are reproduced from main text Figure 5. **B.** Transcript per million (TPM) counts for seven P53 target genes and *TP53* from (A) divide by SMA subgroup. Testing was performed using a one-way ANOVA with multiple-hypothesis correction. **B.** Number of validated P53 targets from P53 target database that are differentially expressed in each subgroup of SMA muscle samples. There is a total of 343 P53 target genes included in this comparison. **D.** Western blot for phosphorylated serine 15 on P53 in SMA and control muscle lysate compared to a vinculin housekeeping control. Quantification is provided on the right. Sample groups were compared using a student’s t-test. **E.** Western blot for GADD45A and vinculin in mouse SMNΔ7 muscle tissue collected at postnatal day 7 (symptom onset) and postnatal day 14 (disease end-stage). Each dot represents an individual mouse sample. Bands are quantified on the right. The distribution of the means was compared using a t-test (not significant) at each age. P . 0.01, * <0.01, ** <0.001, ***<0.0001.

